# Modulation of plant acetyl CoA synthetase activity by post-translational lysine acetylation

**DOI:** 10.1101/2020.01.31.928937

**Authors:** Naazneen Sofeo, Dirk C. Winkelman, Karina Leung, Basil J. Nikolau

**Author notes:** To whom correspondence should be addressed: Basil J. Nikolau: 3254 Molecular Biology Building, Roy J. Carver Department of Biochemistry, Biophysics and Molecular Biology, Iowa State University, Ames, IA 50011-1079, USA. **Authors’ e-mail addresses:** Naazneen Sofeo, Dirk C. Winkelman, Karina Leung, Basil J. Nikolau.

## Abstract

Acetyl-CoA synthetase (ACS) is one of several enzymes that generate the key metabolic intermediate, acetyl-CoA. ACS in plant cells is part of a two-enzyme system that maintains acetate homeostasis, but its post-translational regulation is unknown. In microbes and mammals ACS activity is regulated by the post-translational acetylation of a key lysine residue that is positioned in a conserved domain near the carboxyl-end of the protein sequence. This study demonstrates that the plant ACS activity can be regulated by the acetylation of a specific lysine residue that is homologous to the regulatory lysine residue of microbial and mammalian ACS. The inhibitory effect of the acetylation of residue Lys-622 of the Arabidopsis ACS was demonstrated by site-directed mutagenesis, including the genetic substitution of this residue with the non-canonical N-ε-acetyl-lysine residue. This latter modification lowered the catalytic efficiency of the enzyme by a factor of more than 500-fold. Michaelis-Menten kinetic analysis of the mutant enzyme indicates that this acetylation affects the first half-reaction of the ACS catalyzed reaction, namely the formation of the acetyl adenylate enzyme intermediate. The post-translational acetylation of the plant ACS would thus affect acetate flux in the plastids and overall acetate homeostasis.

**Highlight:** The study demonstrates that acetylation of a specific lysine residue can regulate the activity of acetyl-CoA synthetase, a new regulatory mechanism for maintaining acetate homeostasis.

## INTRODUCTION

The interplay between acetylation and deacetylation of protein lysine residues is a crucial process in regulating many biological processes including chromatin structure, transcriptional regulation and the regulation of cellular metabolism (Bernal *et al*., 2014; Drazic *et al*., 2016; Verdin and Ott, 2015). This post-translational modification mechanism is evolutionarily conserved over a wide range of phyla, and the two enzymes that play a role in this process are: a) lysine acetyl transferases (KATs) that acetylate lysine residues; and b) lysine deacetylases (KDACs) that hydrolyze and remove the acetyl group from the side chain (Drazic *et al*., 2016). In addition, non-enzymatic auto-acetylation of lysine residues occurs at alkaline pH, particularly of lysine residues that are flanked by positively charged amino acids (Narita *et al*., 2019).

Acetyl-CoenzymeA (CoA) synthetase (ACS) is a ∼72 kDa protein that catalyzes the synthesis of acetyl-CoA from acetate and in microbes it plays an important role in fatty acid and polyketide biosynthetic pathways. Moreover, this enzyme generates the substrate that is used in the acetylation of protein lysine residues. In a wide variety of organisms ACS activity is known to be regulated by reversible post-translational acetylation. Specifically, the ACS from *Mycobacterium tuberculosis* can be non-enzymatically auto-acetylated (Li *et al*., 2011). In addition, the enzymatic acetylation of a lysine residue, as a mechanism of ACS regulation, has also been studied in bacterial systems (Bernal *et al*., 2014; Gardner *et al*., 2006; Noy *et al*., 2014). For example, the non-acetylated ACS of *Salmonella enterica* is about 480-fold more active than the acetylated enzyme (Starai *et al*., 2002). The deacetylation of this enzyme is catalyzed by CobB, which is a homolog of the yeast Sir2 deacetylase (Tsang and Escalante-Semerena, 1998). The acetylation of ACS occurs at the side chain of a Lys residue that is positioned in the middle of a conserved structural motif, called A10 (Gulick, 2009). Structural studies of the *S. enterica* ACS indicate that this acetylation does not induce any conformational changes in the enzyme (Gulick *et al*., 2003). However, acetylation significantly affects the ACS first half-reaction, presumably by aligning the acetate-substrate with the active site residue(s) (Gulick *et al*., 2003); this protein acetylation reaction does not affect the second half-reaction catalyzed by *S. enterica* ACS (Starai *et al*., 2002).

In plant systems, such as Arabidopsis, the metabolic function of ACS is not completely clear (Fu *et al*., 2020). Despite the fact that ACS is plastid localized (Lin and Oliver, 2008), unlike other biological systems, ACS is not the physiological source of acetyl-CoA that is the precursor of the plastid-localized *de novo* fatty acid biosynthesis process (Ke *et al*., 2000). Rather, it appears that ACS in plants is important in maintaining acetate homeostasis; acetate being toxic at higher levels of accumulation (Fu *et al*., 2020). In this study, we explored if the acetylation-based regulation of ACS is extrapolatable from bacteria, yeast and algae to affect the activity of this enzyme in plant systems. Specifically, this was explored by a combination of site-specific mutagenesis strategies, including a strategy for expanding the genetic code, and thereby *de novo* reconstruct the post-translational modification of this protein (Chen *et al*., 2018a). The advantage offered by this latter strategy is the ability to generate homogeneous preparations of modified proteins, and thus is a powerful technique to study post-translational modifications (Johnson *et al*., 2010). In this study, we used this strategy to explore the post-translational acetylation of an ε-amine lysine group of the Arabidopsis acetyl-CoA synthetase (atACS) as a potential regulatory mechanism.

## MATERIALS & METHODS

### Generation of atACS variants

The characterization of the atACS ORF was previously described (Behal *et al*., 2002; Ke *et al*., 2000; Lin and Oliver, 2008; Sofeo *et al*., 2019). Incorporating the non-canonical N-ε-acetyl-lysine (Ac-K) residue into the atACS sequence was facilitated by cloning the wild-type ORF sequence into the pCDF-1b vector, which genetically fuses a Hexa-His-tag at the C-terminus of the atACS sequence. The codon encoding for lysine-622 was mutated to the “TAG” stop codon, using Quikchange Lightning Site mutagenesis kit (Agilent Technologies, Santa Clara, CA), and this mutation was confirmed by direct sequencing of the plasmid product. The orthogonal tRNA containing *pTech* plasmid (Venkat *et al*., 2018; Venkat *et al*., 2017) was obtained from Dr. Chenguang Fan (Department of Chemistry & Biochemistry, University of Arkansas) and was co-transformed with the pCDF-1b construct into *E. coli* strain BL21(DE3).

### Expression and purification of wild-type and variant proteins

Wild-type atACS and the lysine-622 mutant variants were expressed and purified as earlier described (Sofeo *et al*., 2019). Namely, the *E. coli* BL21 strains carrying the atACS expression plasmid vector was grown overnight at 37 °C, with agitation at 250 rpm, in 5-10 ml LB medium containing streptomycin (50 μg/ml) and chloramphenicol (50 μg/ml). The overnight culture was used to inoculate 0.25 L LB media containing the same antibiotics. These cultures were grown at 37 °C with agitation at 250 rpm, until the OD_600_ reached ∼0.6. Protein expression was then induced by the addition of IPTG to a final concentration of 0.4 mM. At this time, the culture medium was supplemented with 20 mM nicotinamide to inhibit the endogenous cobB deacetylase (Gallego-Jara *et al*., 2017; Venkat *et al*., 2017), and 2 mM N-ε-acetyl-lysine (Sigma-Aldrich Co., St. Louis, MO), to ensure that this non-canonical amino acid did not limit the expression of the Ac-K atACS variant. The culture was then incubated at 22 °C for 24-48 hours with agitation at 250 rpm. The variant proteins were purified by a process similar to that of the wild-type atACS (Sofeo *et al*., 2019), except all buffers contained 20 mM nicotinamide. The purified atACS proteins were immediately dialyzed into 10 mM HEPES-KOH, pH 7.5, 10 mM KCl, 2mM TCEP, 10% glycerol, concentrated and either characterized immediately or flash frozen in liquid nitrogen and stored at −80°C.

### Gel Filtration Chromatography

Size exclusion gel filtration chromatography was conducted with an AKTA FPLC system (GE Healthcare Life Sciences, Pittsburg, PA) using a Superdex 200 Increase 10/300 GL gel filtration column (GE Healthcare Life Sciences, Pittsburg, PA). A 100 µl aliquot of purified atACS protein (4-10 mg/ml) was injected into the prepacked column, and chromatography was conducted with a buffer consisting of 10 mM HEPES-KOH, pH 7.5, 10 mM KCl, 2 mM TCEP, 10% glycerol, eluted at a rate of 0.4 ml/minute; the eluate was monitored using a UV absorbance detector, at 280 nm.

### Autoacetylation of isolated atACS

The non-enzymatic, autoacetylation of atACS (∼130 µg of purified protein) was conducted in a 1-mL volume of a buffer consisting of 10 mM potassium acetate, 10 mM MgCl_2_, 10 mM ATP, 50 mM Tris-HCl, pH 8.0 (Li *et al*., 2011). Following incubation at 37°C for 2 hours, the protein solution was dialyzed into 10 mM HEPES, pH 7.5, 10 mM KCl, 2 mM TCEP, 10% glycerol.

### Protein analysis

Protein preparations were evaluated by SDS-PAGE, and the acetylation status of atACS was evaluated by Western blot analysis, using an antibody (1:1000 diluted) that reacts with *N*-ε-acetyl-L-lysine (Cell Signaling Technology, #9441). The acetylation status of atACS was also evaluated by mass spectrometric analysis of rLysC (Promega Corporation, Madison, WI) digested protein, using Q Exactive™Hybrid Quadrupole-Orbitrap Mass Spectrometer (Thermo Fisher Scientific Inc., Waltham, MA), housed at the Iowa State University Protein Facility (http://www.protein.iastate.edu). The purified protein preparations of wild-type and Ac-K atACS variants were subjected to SDS-PAGE analysis. Following staining with Coomassie Brilliant Blue, the protein band of interest was excised and digested using an Investigator™ProGest (Genomic Solutions, Digilab Inc, Hopkinton, MA) in 0.5 mL buffer (50 mM Tris, pH 8, 15 mM iodoacetamide and 5 mM DTT) containing 20 µg LysC per mg of purified atACS protein. The peptides were identified using Sequest-HT (Eng *et al*., 1994) as the search engine within Proteome Discoverer (PD) 2.2 (Version 2.2.0.388; Thermo Fisher Scientific). The peptides were analyzed against the ACS protein sequence with settings for four possible missed cleavage sites, with fragment mass tolerance of 0.02 Da precursor mass tolerance of 10 ppm. The possible side-chain modifications that were analyzed include the dynamic acetylation of Lys, static carbamidomethylation of Cys, dynamic deamidation of Asn and Gln and dynamic oxidation of Met residues.

### Spectrophotometric atACS Activity Assay

AtACS activity was measured by coupling the acetate- and CoA-dependent formation of AMP from ATP, to the oxidation of NADH, using the reactions catalyzed by myokinase, lactate dehydrogenase and pyruvate kinase (Sofeo *et al*., 2019). Progress of this reaction was monitored as a decrease in A_340_.

### Circular Dichroism Spectroscopy

CD spectra were obtained with a Jasco J-710 CD spectrometer (JASCO Analytical Instruments, Easton, MD) at the Iowa State University Chemical Instrumentation Facility (https://www.cif.iastate.edu). Spectra were taken at wavelength range of between 190 nm and 260 nm, with the following settings parameter: pitch point 1, speed 20 nm/min, response time 4 sec, bandwidth 1 nm, temperature 20 °C and data mode CD-HT. Spectra where acquired with a 1 mm dichroically neutral quartz cuvette and the average of 3 scans was used for analysis. Acquisition of spectra were obtained with protein samples in 10 mM HEPES-KOH, pH 7.5, 10 mM KCl, 2 mM TCEP and 10% glycerol, and these solutions were diluted into double distilled H_2_O to a final protein concentration of 0.1-0.2 mg/ml. A baseline-spectrum for the same dilution of the buffer was obtained and subtracted from the experimental protein spectrum. Collected spectral data were converted from millidegrees to molar ellipticity ([θ], degrees cm^2^.dmol^-2^). Spectra were analyzed using a suite of algorithms collectively called CDPro (Sreerama and Woody, 2000), utilizing algorithms: SELCON3, CDSSTR, and CONTIN/LL.

## RESULTS

The atACS sequence shares considerable similarity with ACS sequences from a range of diverse biological phyla, and these occur in 10 conserved motifs (A1 to A10) that are distributed evenly throughout the length of these sequences (Gulick, 2009). Three of these motifs (A3, A4 and A5) contain conserved residues that determine the carboxylate substrate specificity of this class of enzymes (Gulick *et al*., 2003; Marahiel *et al*., 1997; Starai and Escalante-Semerena, 2004). Motif A10 contains a highly conserved lysine residue (at position 622 of the Arabidopsis enzyme) (Figure 1). Based on studies of homologous proteins, this lysine residue of the Arabidopsis enzyme may be a target for reversible acetylation, a mechanism that regulates the catalytic capability of these enzymes (Starai *et al*., 2002).

**Figure 1:**
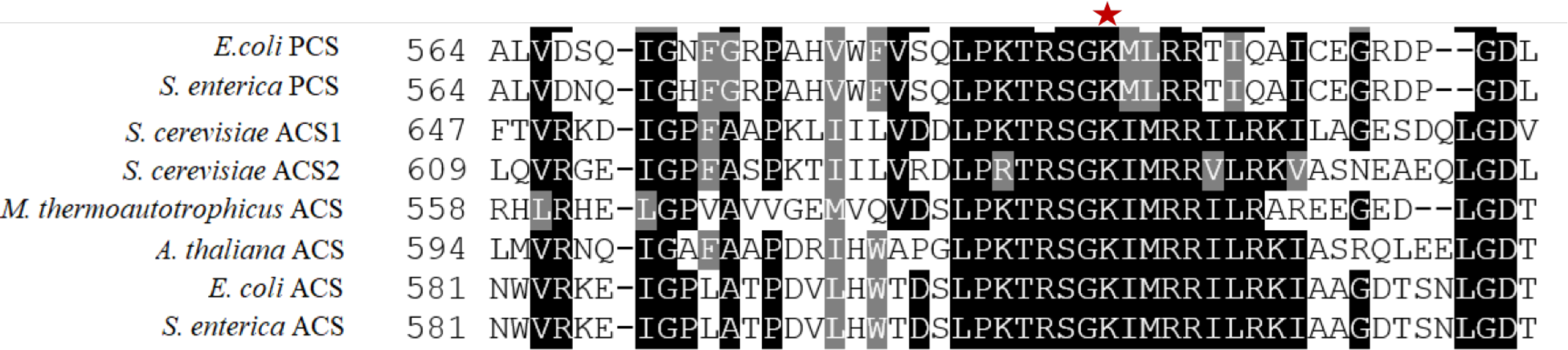
Comparison of the amino acid sequences of motif A10 of the AMP-forming family of acyl-CoA synthetases. Sequence alignments of enzymes that use acetate (ACS) or propionate (PCS) as substrates from different organisms. The red asterisk identifies the conserved lysine residue that is reversibly acetylated. Alignment was performed with Clustal Omega, and white letters on a black background indicate identity, white letters on a gray background indicate similarity, and black letters on a white background indicate no conservation.

Several different experimental strategies were used to evaluate whether acetylation of residue Lys^622^ of atACS modulates catalytic activity of the enzyme. One of these experiments focused on the characterization of mutant variants, in which the Lys^622^ residue was mutated to Ala, Gln or Arg residues. These variant proteins behaved similarly to each other and to the wild-type enzyme in such attributes as: a) stability and yield upon expression in *E. coli*, recovering 10-15 mg of purified protein/L of culture (Figure 2A); b) elution from size exclusion gel filtration chromatography occurs as a single symmetric peak, at an elution volume consistent with the molecular weight of a ∼72-kDa monomer (Figure 2B); c) CD spectra show that all variant proteins are similarly folded as the wild-type enzyme, and secondary structure calculations indicate insignificant differences in the proportion of α-helices, β-sheet, turns and unordered secondary structures among the 3 variant proteins as compared to the wild-type protein (Figure 2C and 2D).

**Figure 2:**
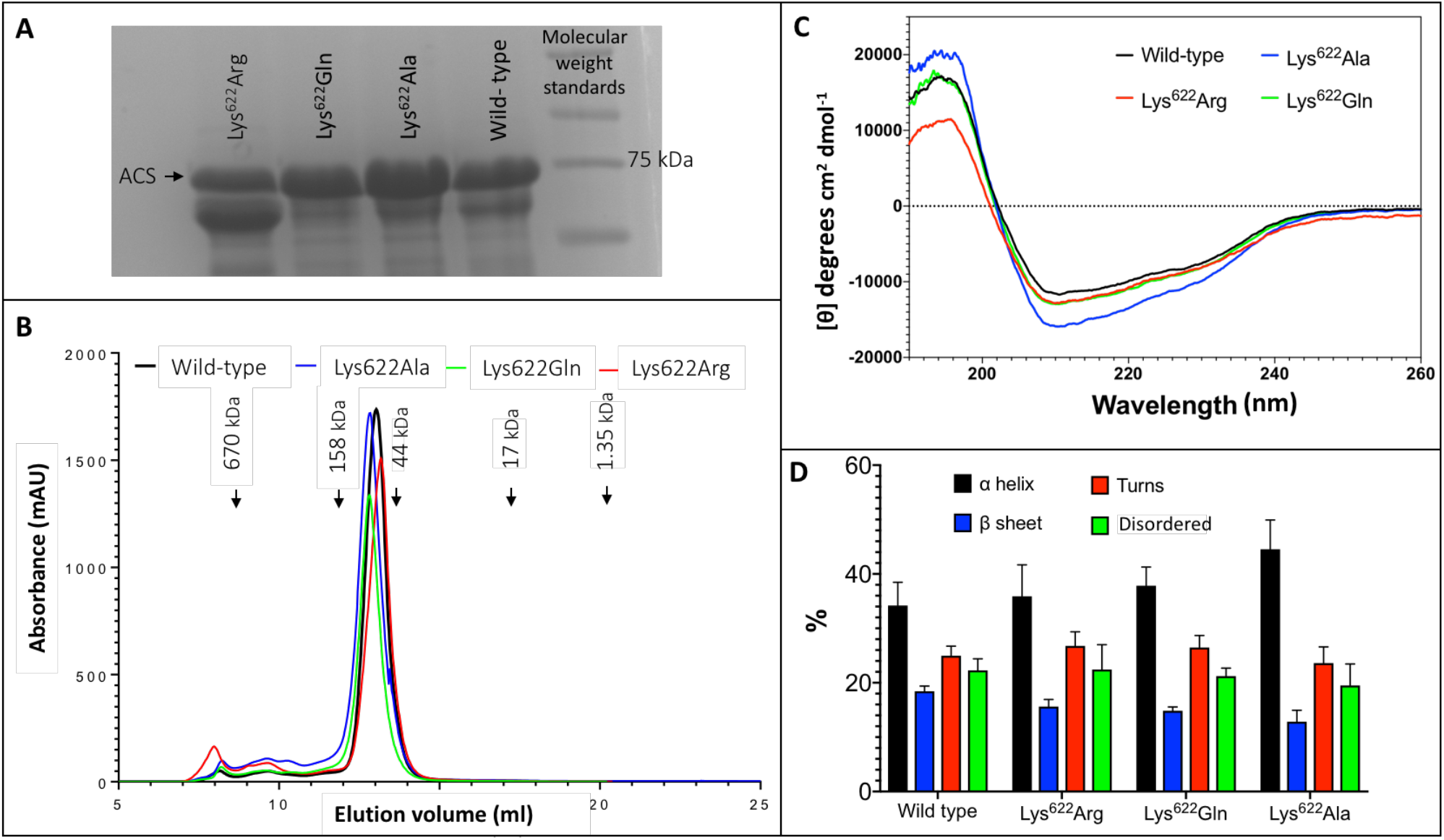
Characterization of ACS Lys^622^ mutants. **A**. SDS-PAGE analysis of purified wild-type and indicated Lys^622^ mutant variants of atACS. **B**. Size exclusion gel filtration chromatography of wild-type and Lys^622^ mutant variants of atACS. **C**. CD spectra of wild-type and Lys^622^ mutant variants of atACS. Measurements are an average of three replicate scans. **D**. Secondary structure composition calculated from the CD spectra of wild-type and Lys^622^ mutant variants of atACS. Errors bars indicate standard errors from 3 replicate CD spectra calculated by 3 algorithms, SELCON3, CDSSTR, and CONTIN/LL.

Enzymatic assays of the three Lys^622^ variants demonstrate that compared to the wild-type enzyme, catalytic competence of these variants is reduced by more than 30-fold. Specifically, the k_cat_ for the Lys^622^Arg mutant is reduced by 30-fold (increasing K_m_ by 25-fold), while the catalytic activity of the Lys^622^Ala and Lys^622^Gln mutants is even further reduced to such low levels that catalytic constants could not be accurately determined (Figure 3A & B).

**Figure 3:**
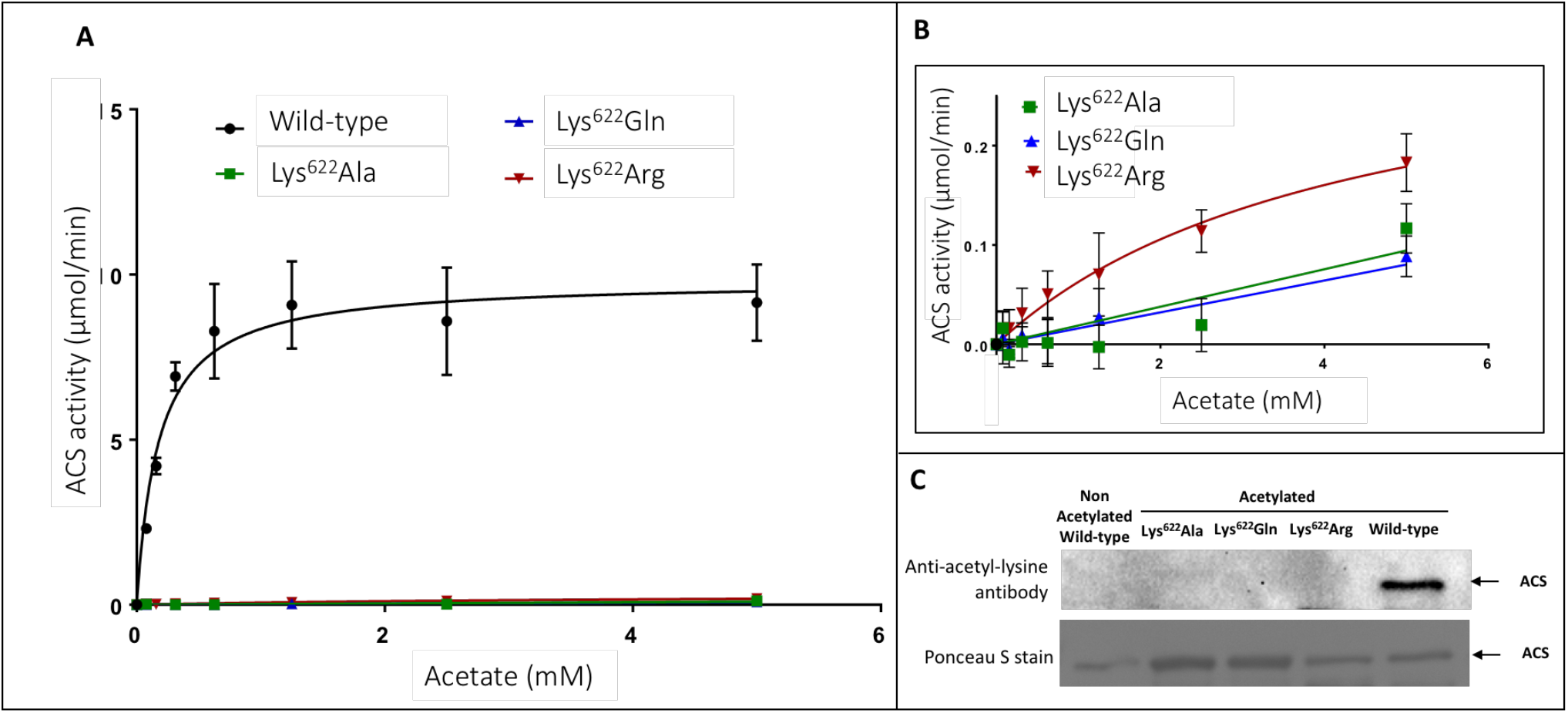
Effect of acetylation on catalytic properties of atACS. **A**. Michaelis-Menten kinetic analysis of atACS Lys^622^ variants. **B**. Expanded view of data presented in Panel A. **C**. Acetylation status of atACS Lys^622^ variants, analyzed by western blot analysis with anti-acetyl-lysine antibody. The top panel is the result from western blot analysis and bottom panel shows Ponceau S staining of the same membrane.

The FPLC purified wild-type and Lys^622^ mutant variants of atACS were chemically autoacetylated *in vitro*, and their acetylation status was determined by western blot analysis using the anti-acetyl-lysine antibody. While the wild-type enzyme is acetylated, as expected all the Lys^622^ variants are not acetylated (Figure 3C). Therefore, these data indicate that this residue may be the site of acetylation of the atACS, which affects enzymatic competence of this enzyme.

Direct assessment of this hypothesis was evaluated by incorporating acetyl-lysine (AcK) specifically at position #622 of atACS. This was accomplished by first engineering a stop codon into the ORF sequence at position-622. The expression of this variant in the absence of AcK results in the expression of a truncated atACS protein that terminates at codon #622. However, when AcK is provided in the medium, because of the expression of a cognate tRNA to read the stop codon as acetyl-lysine (Venkat *et al*., 2018; Venkat *et al*., 2017), AcK is incorporated at position 622, and the resultant atACS-AcK variant is translationally fused at the C-terminus to a His_6_-tag that is part of the expression vector used in these studies. Thus, Ni-column affinity chromatography was used to purify this atACS-AcK-His_6_-tag variant. SDS-PAGE analysis of the recovered protein preparations identified two proteins bands, close to the expected 72-kDa band of atACS (Figure S1. A). Size exclusion gel filtration chromatography of the protein preparations identified two UV-absorbing peaks (A and B) (Figure S1. B). The fractions that encompass these the two protein peaks were pooled separately as indicated (Figure S1. B) and concentrated for further analysis.

These two protein-preparations were subjected to chymotrypsin digestion, and resulting peptides were analyzed by mass-spectrometry to determine their identity. The Peak B derived peptide sequences showed 55% coverage of the atACS sequence, whereas Peak A peptides showed only 4.5% coverage of the atACS sequence (Figure S2 & S3). Additionally, enzymatic assays of Peak A and B preparations indicate that Peak A does not support ACS catalytic activity, whereas Peak B supports the catalysis of acetyl-CoA synthesis (Figure S4). Based on these results therefore, we conclude that Peak A represents a contaminating protein, and that Peak B is the atACS-AcK variant, and all further characterizations were conducted by pooling fractions that encompassed Peak B.

Direct demonstration that this isolated atACS-AcK variant is acetylated at the targeted Lys^622^ residue was obtained by mass-spectrometric analysis of the LysC-digested protein, and these data were compared to the mass-spectrometric analysis of the LysC digested wild-type atACS protein. The sequence coverage from these analyses were 40% for the wild-type atACS protein, whereas coverage of the atACS-Ack variant was ∼62% (data not shown). Although the lysine-622 containing peptide could not be identified in the LysC digest of the wild-type atACS, it was successfully detected from the atACS-Ack variant protein. The acetylated lysine-622 peptide sequence was identified as, TRSGK^Ac^IMRRILRK. This was based on the monoisotopic m/z value of 415.01038 Da (+0.76 mmu/+1.84 ppm), with a charge of +4 and theoretical mass of the MH+ ion of 1657.01967 Da (Figure 4).

**Figure 4:**
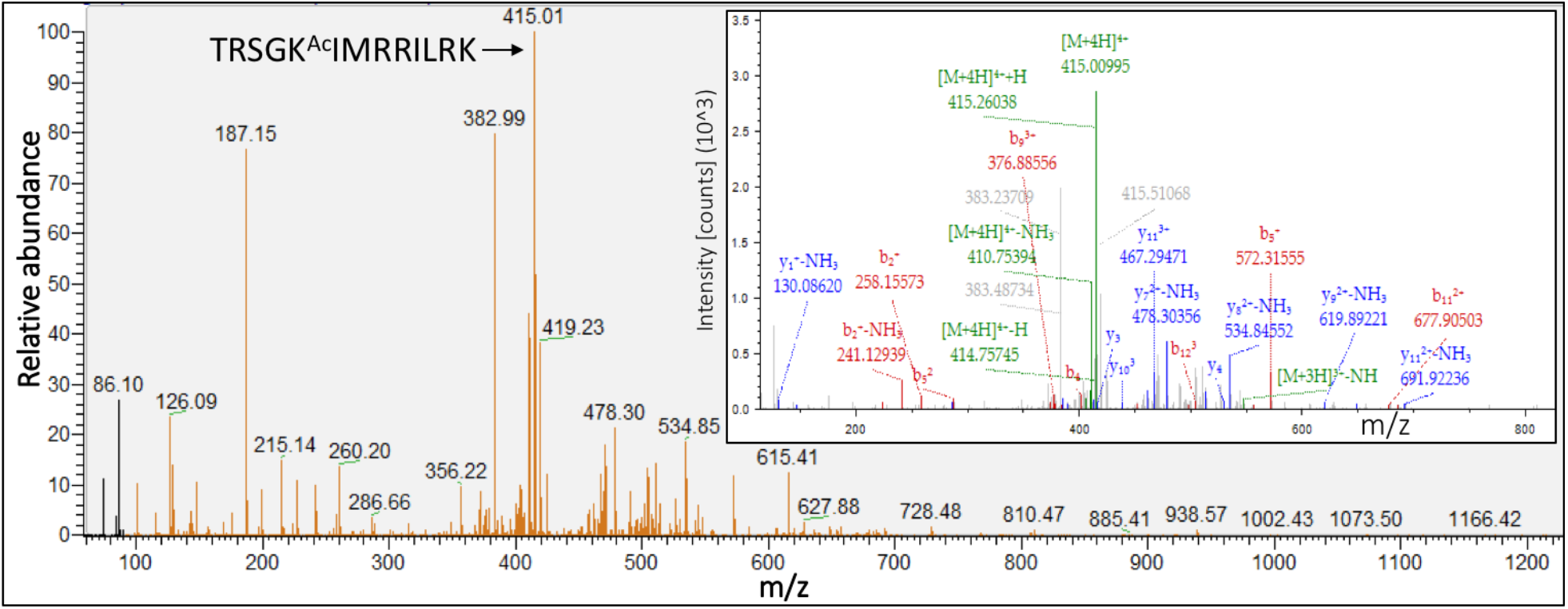
Mass spectrometric identification of acetylated Lys-622 containing peptide. LC-MS/MS spectra and MS2 fragmentation pattern (inset) of the LysC generated peptide containing lysine 622 from atACS Ac-K variant.

Comparing the Michaelis-Menten kinetic parameters (K_m_ and k_cat_) of atACS to the atACS-Ack variant indicates that acetylation of the enzyme affects its ability to catalyze acetyl-CoA synthesis (Figure 5). Specifically, as compared to the wild-type atACS, the atACS-Ack variant shows a 34-fold decrease in k_cat_, and a 15-fold increase in K_m_ for the acetate substrate (Figure 5).

**Figure 5:**
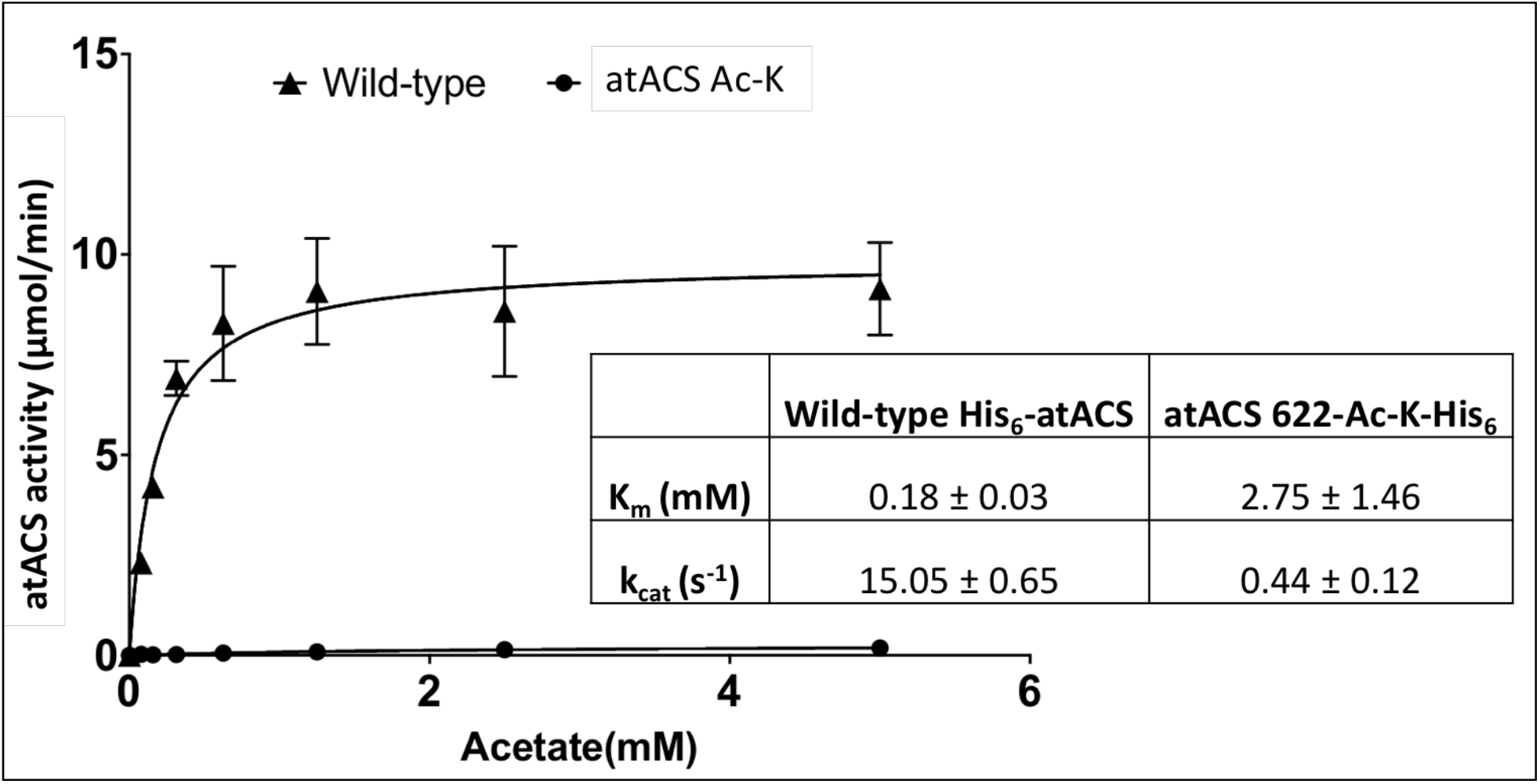
Catalytic capabilities of wild-type atACS (▴) and atACS Ac-K variant (•). (Inset table) Kinetic parameters of wild-type atACS and atACS Ac-K variant.

## DISCUSSION

It has become increasingly apparent that the post-translational acetylation-deacetylation cycle of the ε-amino group of lysine residues is an important mechanism for regulating many biological processes (Chen *et al*., 2018a; Christensen *et al*., 2019; Hossain and Tsang, 2019; Hu *et al*., 2019; Wang and Wang, 2019). This modification reaction is one of the broader suites of mechanisms by which the chemo-physical properties of side chains of amino acid residues can be diversified from the 20 genetically encoded canonical amino acids. Moreover, because these modifications are often reversible, and enzyme catalyzed, these reactions provide a rapid mechanism for regulating biological processes. Specifically, one can rationalize that acetylation of the ε-amino group of a lysine residue changes the character of that side chain from a highly polar, positively charged residue to a non-polar side chain that is capable of participating in weaker H-bonding interactions as both a H-bond donor or H-bond acceptor. Such drastic changes in the properties of such a side chain can generate molecular forces that confer structural alterations in proteins, which thereby render altered functionality.

Because ACS has a role in generating the substrate for such acetylation modification reactions, the post-translational acetylation-deacetylation of this enzyme has potential to generate an autoregulatory loop that could enhance or dampen such controlling mechanisms. Indeed, in microbes and mammals ACS catalytic activity can be regulated by the acetylation of a specific lysine residue that occurs in a conserved region of these enzymes (Gardner *et al*., 2006; Hallows *et al*., 2006; Noy *et al*., 2014; Schwer *et al*., 2006; Starai *et al*., 2003). For example, acetylation of a specific lysine residue in the A10 motif of the bacterial ACS inactivates catalysis (Gardner *et al*., 2006; Noy *et al*., 2014). Similarly, with the mammalian enzyme, acetylation completely deactivates catalysis, and deacetylation by sirtuin reactivates enzymatic activity (Hallows *et al*., 2006; Schwer *et al*., 2006). In contrast, acetylation of the analogous lysine residue in homologous long chain acyl-CoA synthetases increases catalytic activity (Chen *et al*., 2018b).

Moreover, global proteomics studies have identified that the acetylation of this specific Lys residue occurs *in vivo* in many bacteria, fungi and animals (Desiere *et al*., 2006; Gulick *et al*., 2003), including the well characterized *E. coli* and *S. enterica* ACS. Unfortunately, the Lys-622 residue that was the subject of this study is in a region of the atACS protein that contains a plethora of Lys and Arg residues, which has prevented it from being detected in global proteomics studies that typically utilize trypsin as the protease to generate peptides for MS analysis. In this study therefore, a series of experiments were conducted that expanded upon prior studies and demonstrated by a combination of different strategies that the plastid localized plant ACS is susceptible to post-translational acetylation mechanism, which can modulate catalytic capability. Specifically, our western blot and mass spectrometric studies demonstrate that acetylation of Lys-622 of atACS reduces catalytic efficiency (k_cat_/K_m_) of the enzyme by ∼522-fold. The importance of Lys-622 acetylation to ACS catalysis was demonstrated by a number of mutagenesis experiments. Specifically, eliminating this side chain, as in the Lys622Ala variant eliminates catalysis. Moreover, the conservative substitution of this residue (i.e., Lys622Arg variant), which maintains the positive side chain characteristic also eliminates ACS catalysis. Furthermore, the substitution of Lys-622 with a residue that is capable of H-bond formation (i.e., Lys622Gln variant) has the same inactivation result. These mutations do not affect the overall gross folding, oligomeric state, or the *in vivo* stability of the recombinant atACS protein, even though it renders the enzyme inactive. Hence, these results indicate the importance of this lysine residue in supporting the catalytic mechanism of this enzyme, but do not directly address whether this Lys residue is acetylated.

We therefore used an alternative strategy to address this question, namely using an already acetylated lysine residue that was genetically inserted into the protein sequence at the target position during the *in vivo* synthesis of the atACS protein. Such a strategy has been used to not only incorporate N-ε-acetyl-lysine at targeted sites, but also a number of different non-canonical amino acids (Johnson *et al*., 2010). The strategy uses an orthogonal pair of a suppressor tRNA and its respective amino acyl tRNA synthetase that recognizes a specific non-canonical amino acid, and that amino acid is genetically programmed to be incorporated during the translation process of a host, typically *Escherichia coli* (Chen et al., 2018a). The position for incorporating the non-canonical amino acid is defined by stop-codon mutagenesis of the protein-coding ORF, which is recognized by the anticodon of the suppressor tRNA that carries the non-canonical amino acid (Xie and Schultz, 2006). Using this strategy we generated homogeneous preparations of modified protein (i.e., atACS-Ack) for characterization. Comparing the kinetic parameters of the wild-type and atACS-Ack variant, we found that the acetylated enzyme showed an increase in the K_m_ and decrease in the k_cat_ values for the acetate substrate. Collectively therefore, these results establish that acetylation of Lys-622 of atACS results in the inactivation of its catalytic capability. Hence, reversible acetylation of this residue would be an important mechanism by which the activity of this enzyme is modulated *in vivo*, especially to accommodate changes in cellular acetate and/or acetyl-CoA levels.

Consistent with its role in catalysis, the experimentally determined structure of the acetylated and non-acetylated bacterial ACS (Fox III *et al*., 2017; Gulick *et al*., 2003) has established that this Lys residue is near the active site pocket during the first half of the reaction, when acetate and ATP bind and react to form the acetyl-adenylate intermediate and release the pyrophosphate product (Gulick, 2009; Gulick *et al*., 2003). MStructural modeling of the atACS indicates a similar configuration for Lys-622, which indicates that acetylation inhibits the first half-reaction catalyzed by atACS.

The *in vivo* physiological role of ACS in plants was initially thought to be the enzymatic supplier of the acetyl-CoA substrate required for *de novo* fatty acid biosynthesis in plastids (Behal *et al*., 2002). However, genetic and biochemical characterizations established that ACS is not needed for this metabolic role (Ke *et al*., 2000), and more recent genetic characterizations establish that ACS is part of a 2-enzyme system that plants use to maintain acetate homeostasis (Fu *et al*., 2020). Because acetyl-CoA is a crucial intermediate of metabolism that juxtaposes anabolic and catabolic processes, and is also a critical component of many regulatory processes associated with acetylation of controlling components (e.g., histone acetylation or N-terminal and/or amino acid side-chain acetylation), one can envision that its generation is highly regulated. Hence, the post-translational acetylation mechanism identified herein for modulating atACS activity could provide plants with the ability to rapidly adapt to changing environmental and developmental conditions and maintain cellular acetyl-CoA and acetate homeostasis.

## Supplementary data

The following supplementary data are available at JXB online.

**Figure S1**: Purification of atACS-AcK variant.

**Figure S2**: Mapping of peptides (green shaded residues) identified by mass-spectrometric analysis of chymotryptic digest of Peak A protein, to the atACS sequence.

**Figure S3**: Mapping of peptides (green shaded residues) identified by mass-spectrometric analysis of chymotryptic digest of Peak B protein, to the atACS sequence.

**Figure S4**: Acetate dependence of the ACS activity of the FPLC purified atACS-AcK variant.

## Acknowledgement

This work was partially supported by the State of Iowa, through the Center of Metabolic Biology, and by the National Science Foundation (Award No. EEC-0813570), which supported the Engineering Research Center for Biorenewable Chemicals (CBiRC; www.cbirc.iastate.edu). Karina Leung acknowledges support from the Research Experience for Undergraduates program of the National Science Foundation, provided through CBiRC. The authors thank Dr. Chenguang Fan of Department of Chemistry & Biochemistry, University of Arkansas for gifting the *pTech* vector used for expressing orthogonal tRNA synthetase machinery, and Iowa State University Protein Facility (specifically Joel Nott) for providing assistance and guidance with protein mass spectrometric analysis.

## Author contributions

RG-C: Conceptualization; NS, BJN, and RG-C: data curation; NS, DCW, and RG-C: formal analysis; NS, DCW, and RG-C: funding acquisition; BJN, and RG-C: investigation; NS, DCW, KL, and RG-C: methodology; NS, DCW, and RG-C: project administration; BJN, and RG-C: resources; BJN, and RG-C: supervision; NS, BJN, and RG-C: Visualization; NS, BJN, and RG-C: writing – original draft preparation; NS, DCW, and RG-C: writing – review & editing; BJN.

