## Supplemental Figures for "Modulation of plant acetyl CoA synthetase activity by post-translational lysine acetylation"

**Supplementary Figures**
Sofeo et al.

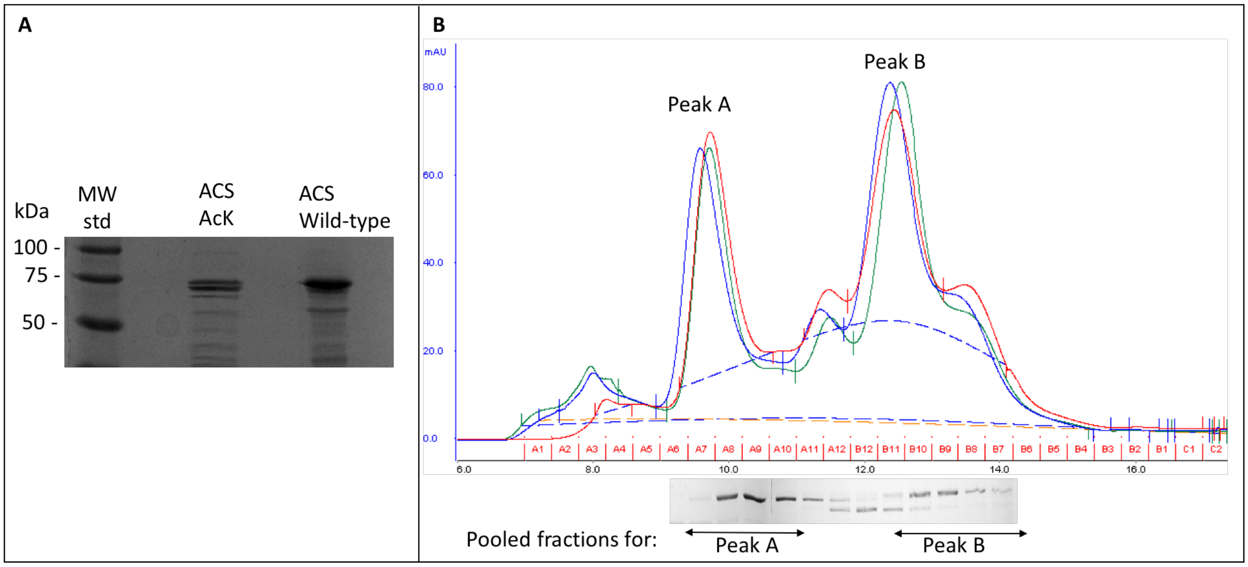


**Figure S1**: Purification of atACS-AcK variant. **A**. Coomassie stained SDS-PAGE analysis of the purified protein preparations. Wild-type atACS and atACS-AcK carried a $His_6$-tag at N-terminus and C-terminus, respectively. **B**. Size exclusion gel filtration chromatography of three preparations of atACS-Ack variants. Aliquots of each peak; Peak A, fraction A7-A10 and Peak B, fraction B10-B7, were analyzed by SDS-PAGE and stained with Coomassie Brilliant Blue (shown under the graph). The indicated fractions were pooled and subjected to further analysis.

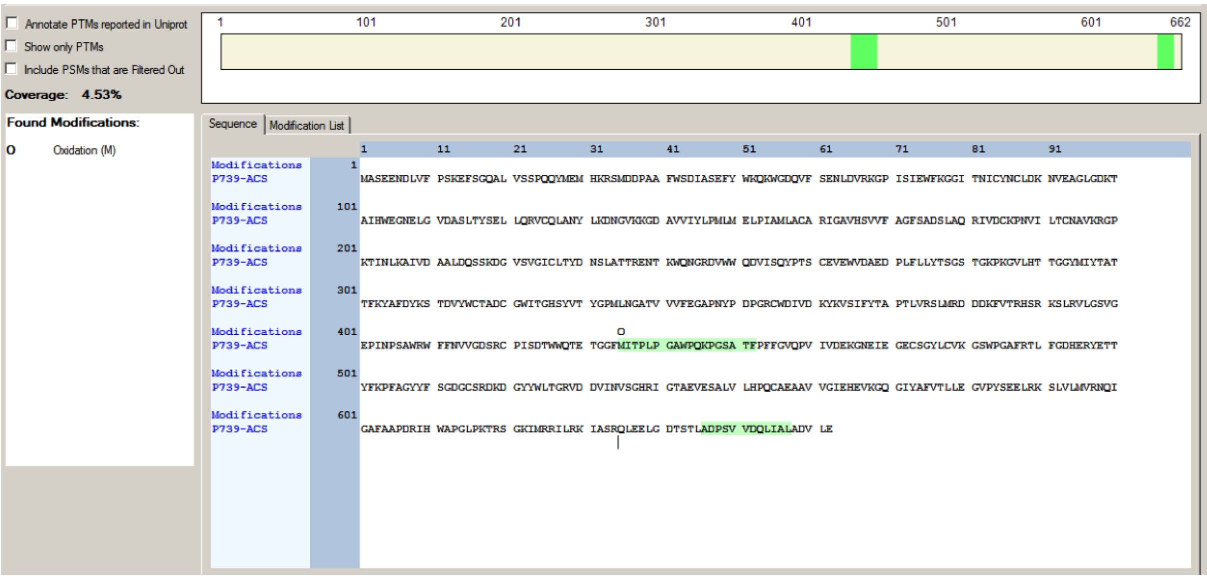

**Figure S2**: Mapping of peptides (green shaded residues) identified by mass-spectrometric analysis of chymotryptic digest of Peak A protein, to the atACS sequence; coverage is 4.5% of the atACS sequence.

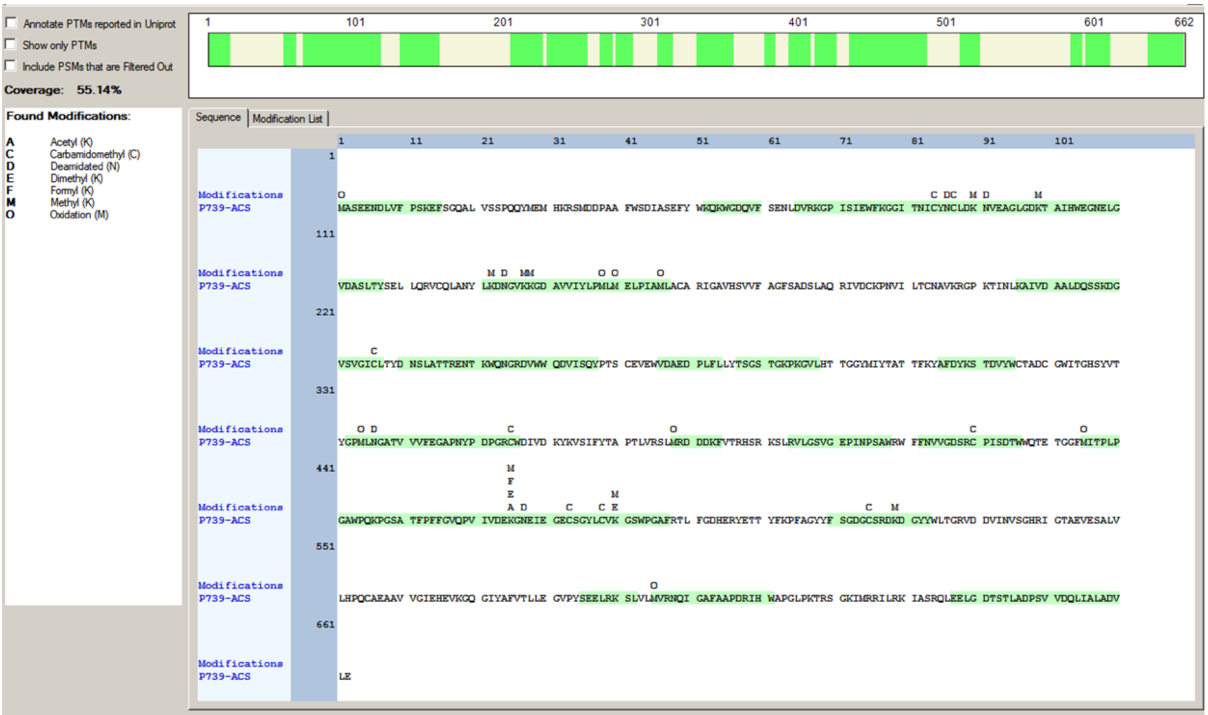

**Figure S3**: Mapping of peptides (green shaded residues) identified by mass-spectrometric analysis of chymotryptic digest of Peak B protein, to the atACS sequence; coverage is 55% of the atACS sequence.

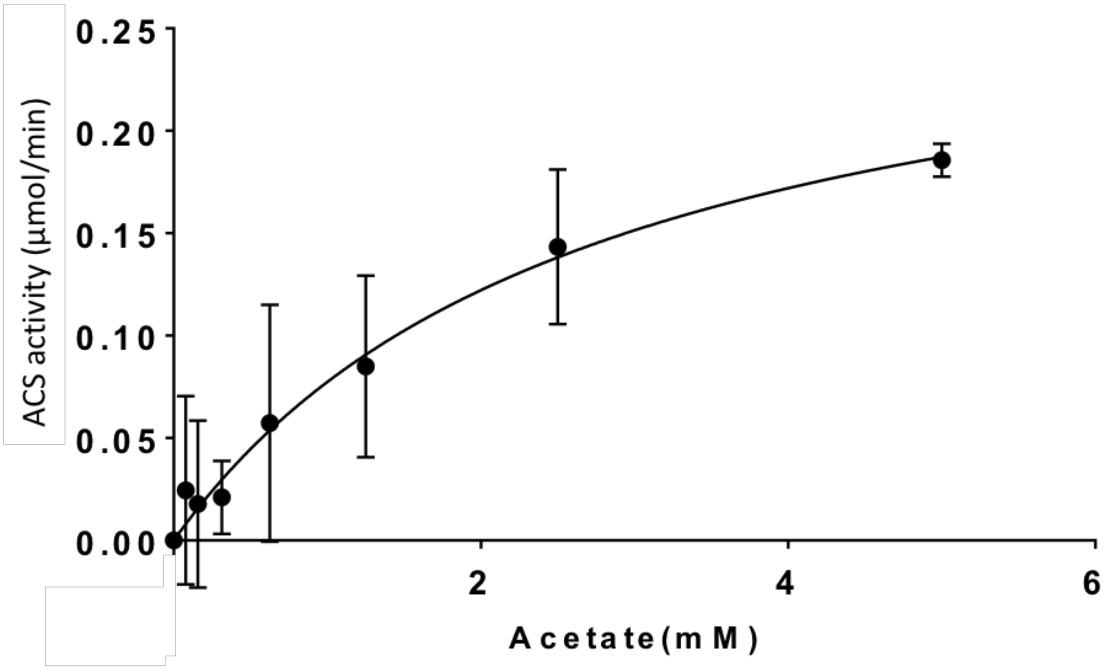


**Figure S4**: Acetate dependence of the ACS activity of the FPLC purified atACS-AcK variant (Peak B). Peak A did not show any detectable activity.
